# SPIKE-TRIGGERED INTRASPINAL MICROSTIMULATION IMPROVES MOTOR PERFORMANCE IN AN AMBULATORY RAT MODEL OF SPINAL CORD INJURY

**DOI:** 10.1101/2021.06.21.449323

**Authors:** Jordan A. Borrell, Domenico Gattozzi, Dora Krizsan-Agbas, Matthew W. Jaeschke, Randolph J. Nudo, Shawn B. Frost

**Author notes:** Corresponding Author: Shawn B. Frost, PhD, Dept. Rehabilitation Medicine, University of Kansas Medical Center. Mail Stop: 1059, 3901 Rainbow Blvd., Kansas City, KS 66160.

## Abstract

The purpose of this study was to determine if spike-triggered intraspinal microstimulation (ISMS) results in improved motor performance in an ambulatory rat model of spinal cord injury (SCI). Experiments were carried out in adult male Sprague Dawley rats with 175 kdyn moderate T8 contusion injury. Rats were randomly assigned to one of two groups: Control or Activity Dependent Stimulation (ADS) therapy. Four weeks post-SCI, all rats were implanted with a recording electrode in the left hindlimb motor cortex and a fine-wire, custom-made stimulating electrode in the contralateral lumbar spinal cord. Intracortical and intraspinal microstimulation were used to find sites of similar hip representation areas, which were paired together for ADS therapy. In the ADS therapy group, spike-stimulus conditioning was administered for 4 hours/day, 4 days/week, for 4 weeks via a tethered cable in a testing chamber. During therapy sessions, single-unit spikes were discriminated in real time in the hindlimb motor cortex and used to trigger stimulation in the spinal cord ventral horn. The optimal stimulus intensity (50% ISMS movement threshold) and spike-stimulus delay (10ms) determined in preliminary anesthetized preparations were used during ADS. Control rats were similarly implanted with electrodes but did not receive stimulation therapy. Motor performances of each rat were evaluated before SCI contusion, once a week post-SCI for four weeks (prior to electrode implantation), and once a week post-conditioning for four weeks. Behavioral testing included BBB scoring, Ledged Beam walking, Horizontal Ladder walking, treadmill kinematics via the DigiGait and TreadScan system, and open field walking using OptiTrack kinematic analysis. BBB scores were significantly improved in ADS rats compared to Control rats after 1 week of therapy. In the ADS therapy rats, BBB scores were significantly improved after two weeks of ADS therapy when compared to pre-therapy. Foot fault scores on the Horizontal Ladder were significantly lower in ADS rats compared to pre-therapy ADS and Control rats after 1 week of therapy and returned to pre-injury measures after three weeks of ADS therapy. The Ledged Beam test and kinematic analysis using the DigiGait and TreadScan system showed deficits after SCI in both ADS and Control rats but there were no significant differences between groups after 4 weeks of ADS therapy. These results show that activity dependent stimulation after spinal cord injury using spike-triggered ISMS enhances behavioral recovery of locomotor function as measured by the BBB score and the Horizontal Ladder task.

## Introduction

Injury to the central nervous system causes a disconnect between distant populations of neurons which results in chronic motor deficits. As a result, impairment severity is related to the function of remaining viable neurons. By focusing on remaining neurons and their pathways after injury, a primary goal of current researchers has been to leverage the nervous system’s intrinsic capacity for reorganization. One innovative device-based approach utilizes activity-dependent stimulation (ADS) paradigms that record and digitize extracellular neural activity from an implanted microelectrode, discriminate individual action potentials in real time, and deliver small amounts of electrical current to another microelectrode implanted in a distal population of neurons (Guggenmos et al., 2013; Nishimura et al., 2013b; Zimmermann and Jackson, 2014; McPherson et al., 2015; Capogrosso et al., 2016). This approach is based on mechanisms underlying neuroplasticity (Hebb, 1949; Jackson and Zimmermann, 2012). ADS paradigms aim to strengthen the remaining pathways after injury by inducing neuroplasticity to restore or improve lost motor function.

A spinal cord injury (SCI) causes damage to descending motor pathways by disrupting the signals from the brain to the spinal cord and nerve roots at and distal to the level of injury. This lack of input from the brain to neurons below the injury in the spinal cord can result in some level of paralysis. However, even without input from the brain, studies have shown that circuits located below the injury, specifically in the lumbar segments of mammals, can fabricate coordinated patterns of leg motor activity (Sherrington, 1892; Kiehn, 2006). As a result, researchers have developed various neuromodulatory approaches to reconnect brain signals to the spinal cord pattern generators by “closing the loop” with brain-machine interfaces (Zimmermann and Jackson, 2014). Recent studies using closed-loop interfaces between the brain and spinal cord have demonstrated improvement in locomotion (Capogrosso et al., 2016) and functional upper limb movement (Nishimura et al., 2013a) after spinal cord lesions; potentially via strengthened connections or enhanced plasticity in descending motor pathways.

These efforts encouraged us to develop an understanding of the connection between the brain and spinal cord after a bilateral thoracic spinal cord contusion that mimics those seen in clinical spinal cord injuries (Krizsan-Agbas et al., 2014; Frost et al., 2015; Borrell et al., 2020a; Borrell et al., 2020b). Through using SCI models, we can create neuromodulation protocols that can more easily translate into clinical use for rehabilitation. This framework guided the design of a brain-machine-spinal cord interface (BMSI) that uses activity dependent stimulation (ADS) to strengthen motor pathways that may still be intact after SCI (Shahdoost et al., 2016). ADS administered in a closed-loop manner uses neural activity to trigger stimulation. In our ADS paradigm, an implanted recording microelectrode in the hindlimb motor cortex records and digitizes extracellular neural activity, discriminates individual action potentials in real time, and delivers small amounts of electrical current to another stimulating microelectrode implanted in the ventral horn of the lumbar spinal cord distal to the level of the injury.

The ability of the motor cortex to evoke spinal cord activity below the level of an injury demonstrates that some descending motor pathways to the lumbar cord remain intact after a moderate spinal cord contusion (Borrell et al., 2020a). Furthermore, the administration of ADS between the hindlimb motor cortex and hindlimb spinal cord has shown that ADS causes a plastic change in the intact pathways of rats with a thoracic contusion to the spinal cord (Borrell et al., 2020b). The current study was conducted to determine if application of ADS therapy (i.e., spike-triggered intraspinal microstimulation) results in improved motor performance in an ambulatory rat model of SCI. The goal of this rehabilitation intervention is to facilitate and direct intrinsic synaptic plasticity in spared motor pathways and circuits that would result in improved motor performance.

## Materials and Methods

### General Experimental Design

To determine if spike-triggered intraspinal microstimulation (ISMS) therapy results in improved motor performance in an ambulatory rat model of spinal cord injury (SCI), extracellular neuronal activity was recorded from the hindlimb motor cortex and used to trigger ISMS in the ventral horn of the lumbar spinal cord (Figure 1). This procedure is hereafter called activity-dependent stimulation (ADS). Motor performance of rats receiving ADS therapy was scored on five behavioral tasks at pre-specified time points throughout the study. Behavioral scores were analyzed and compared to pre-therapy scores and scores of a non-stimulation control group to examine for improvement in motor performance.

**Figure 1.**
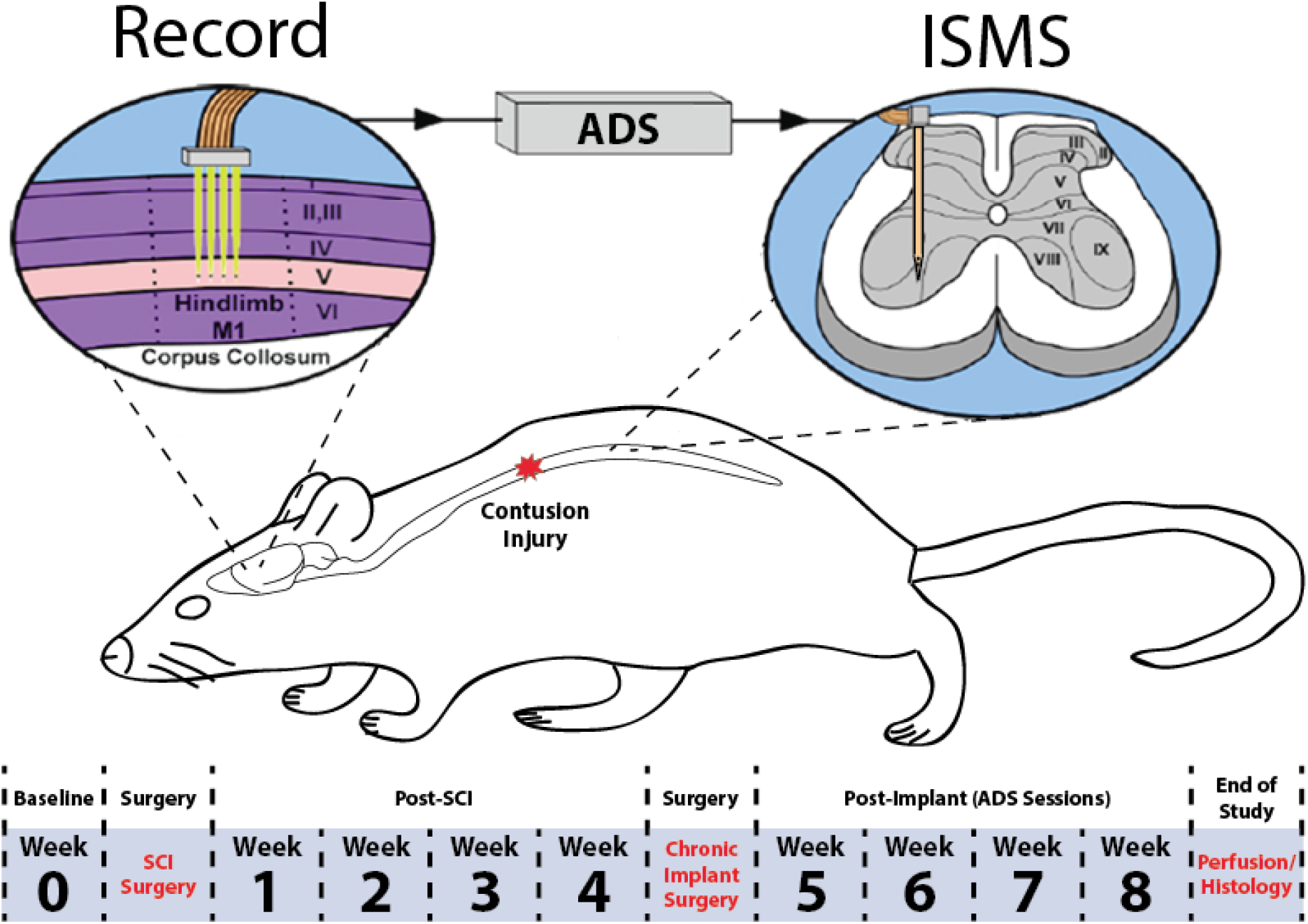
Overview of experimental design. A 175 kDyn spinal cord contusion was administered at the T8 vertebral segment. Four weeks after injury, rats were implanted with a recording electrode in the hindlimb motor cortex and a fine wire electrode in the ventral horn of the lumbar spinal cord. Behavioral tests were conducted before (i.e., baseline) SCI and once per week after each surgery for the duration of the study. Activity dependent stimulation was administered for four weeks post-implant in the ADS group. No stimulation was administered in the control group. ADS sessions occurred during weeks 5-8 for 4 hours/day and 4 days/week.

### Subjects

Twenty adult male Sprague Dawley rats were selected for this study. At the beginning of the study, body weights ranged from 305 to 416 g (mean ± SE = 360.21 ± 9.23 g). Ages ranged from 64 to 106 days old (mean ± SE = 82.89 ± 3.36 days old). Rats were randomly assigned to one of two groups: ADS group and Control group. Each group underwent the same surgical procedures and behavioral analysis described below. Seven rats were removed from the study due to intraoperative complications (SCI surgery, n = 3; implant surgery, n = 4) while one rat was removed for failing to perform the behavioral tasks (n = 1), resulting in a final complement of six rats in the ADS Group (n = 6) and six rats in the Control Group (n = 6). This study was performed in accordance with all standards in *Guide for the Care and Use of Laboratory Animals* (Institute for Laboratory Animal Research, National Research Council, Washington, DC: National Academy Press, 1996). The protocol was approved by the University of Kansas Medical Center Institutional Animal Care and Use Committee.

### Experimental Timeline

The experimental timeline is displayed in Figure 1. Each of the behavioral tests was conducted once per week throughout the duration of the study to evaluate motor performance pre-SCI (i.e., baseline), for 4 weeks post-SCI (i.e., prior to electrode implantation and ADS conditioning), and 4 weeks post-ADS conditioning (i.e., after each week of ADS conditioning). First, rats were acclimated to each of the behavioral tasks. Then, baseline performance was assessed. One to three days after baseline assessment, a surgical procedure was conducted to produce a contusive injury at the T8 vertebral segment. During weeks 1-4 post-SCI, behavioral assessment was conducted once per week to evaluate the effects of the SCI on motor performance and the extent of any spontaneous recovery. Then, a second surgical procedure was conducted to implant a recording electrode in the hindlimb motor cortex and a fine wire electrode in the ventral horn of the lumbar spinal cord. After implants, ADS therapy sessions were conducted during weeks 5-8. During the ADS therapy period, behavioral assessment was conducted once per week at the end of the week, i.e., after each of the four ADS sessions. At the end of the 8-week study, the rats were humanely euthanized, and the spinal cords assessed for the extent of the injury.

### General Surgical Procedures

Both surgical procedures (SCI surgery; chronic implant surgery) followed the same general protocol. After an initial stable anesthetic state was established using isoflurane anesthesia and the scalp and back shaved, isoflurane was withdrawn, and an initial dose of ketamine hydrochloride (100 mg /kg IP)/xylazine (5 mg /kg IP) was administered. The anesthetic state was maintained with subsequent doses of ketamine (10 mg IP or IM) and monitored via pinch and corneal reflex responses. Additional doses of ketamine were administered if the rat reacted to a pinch of the forepaw/hindpaw, had a positive corneal reflex, or exhibited increased respiration rate. At the conclusion of the surgery, 0.25% bupivacaine HCl was applied locally to the skin incision site. Buprenex (0.01 mg/kg, S.C.) was administered immediately after surgery and 1 day later. All animals were monitored daily until the end of the experiment. Before and after surgery, rats received a S.C. injection of 30,000 U of penicillin (Combi-Pen 48).

### Spinal Cord Contusion

Spinal cord contusion procedures followed those described in our previous studies (Krizsan-Agbas et al., 2014; Borrell et al., 2020a; Borrell et al., 2020b). Animals underwent a T8 laminectomy and 175 kDyn moderate impact contusion injury using an Infinite Horizon spinal cord impactor (Precision Systems and Instrumentation, LLC, Fairfax Station, VA). Displacement distance reported by the impactor software for each contusion was recorded at the time of surgery and was used as an initial quantitative marker for successful impact. Starting the first day after surgery, daily penicillin injections were given in 5 mL saline throughout the first week to prevent infections and dehydration. Bladders were expressed twice daily until animals recovered urinary reflexes. From the second week onward, animals were supplemented with vitamin C pellets (BioServ, Frenchtown, NJ) to avert urinary tract infection.

### Chronic Electrode Implantation

Rats were placed in a Kopf small-animal stereotaxic frame (David Kopf Instruments^®^, Tujunga, CA) and the incisor bar was adjusted until the heights of lambda and bregma were equal (flat skull position). The cisterna magna was punctured at the base of the skull to reduce edema during mapping and implantation. Before the craniectomy, five titanium screws were placed at various positions on the skull as shown in Figure 2. These screws served as anchors for a head cap made of dental acrylic that encased the chronic recording array, headstage, and supporting hardware on the skull.

**Figure 2.**
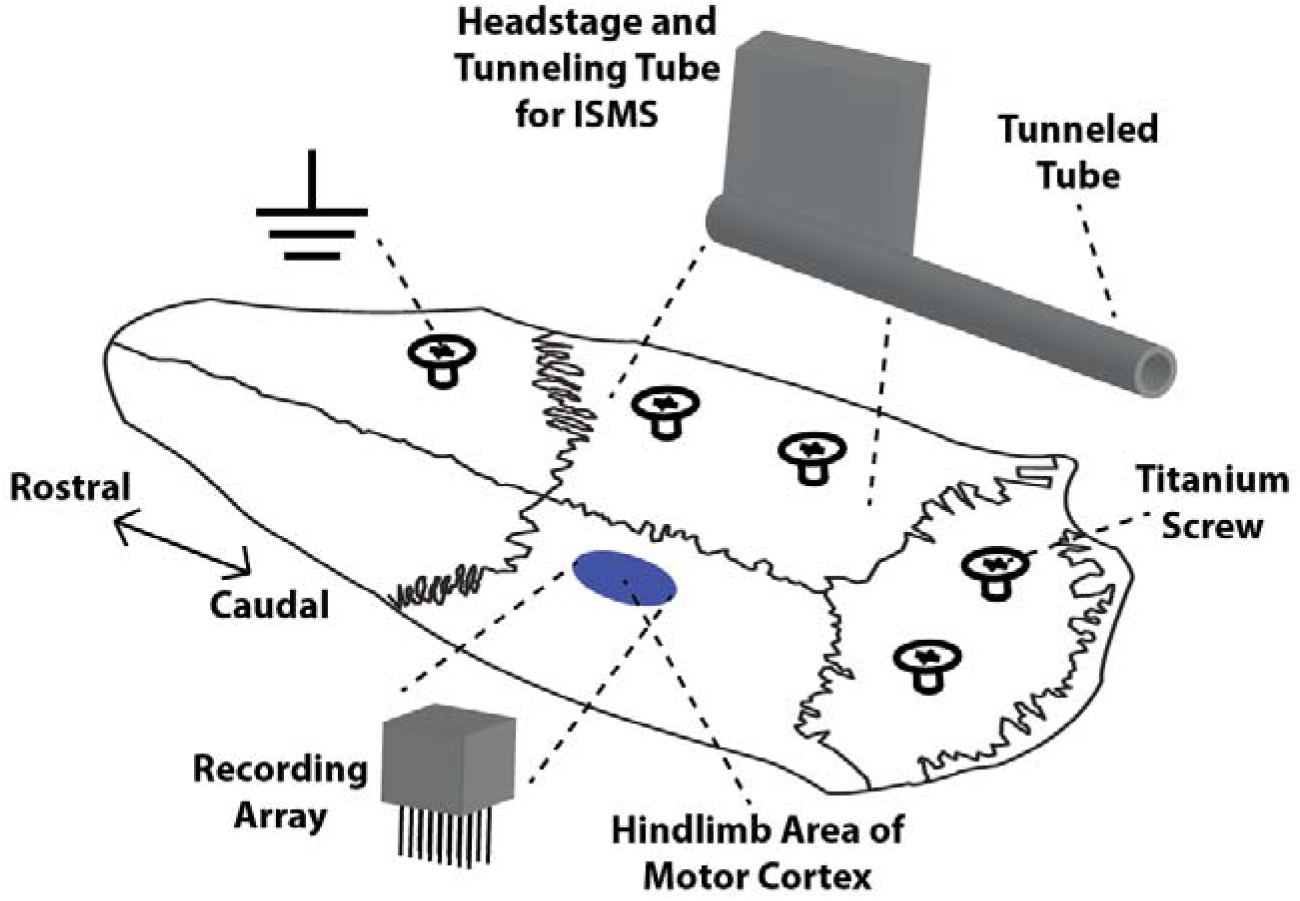
Placement of chronic microelectrode array, headstage, and supporting hardware on the skull of a rat. Dental acrylic was used to build a protective cap and anchor the array and headstage to the skull via five titanium screws. The recording array was implanted in the left hindlimb motor cortex. The headstage for the stimulating electrode was mounted on the right side of the skull while the microwires were tunneled via a tunneling tube to the lumbar spinal cord. The ground wires for both electrodes were wrapped around the rostral-most titanium screw.

For placement of the recording array in the cortex, a craniectomy was performed over the hindlimb area (HLA) of motor cortex in the left hemisphere. The location of the craniectomy was guided by previous rat cortical motor mapping studies (Frost et al., 2013; Frost et al., 2015). After the dura over the cranial opening was incised, a small motor map was derived using intracortical microstimulation (ICMS) using techniques similar to those in our previous studies (Frost et al., 2013; Frost et al., 2015). Since hindlimb movements cannot be evoked using ICMS after the SCI, the location of the hip representation of HLA was inferred by proximity to surrounding representations. Based on our prior ICMS studies, the hip representation is bordered laterally by the trunk area and caudally by the forelimb area of motor cortex (Figure 3 Left).

**Figure 3.**
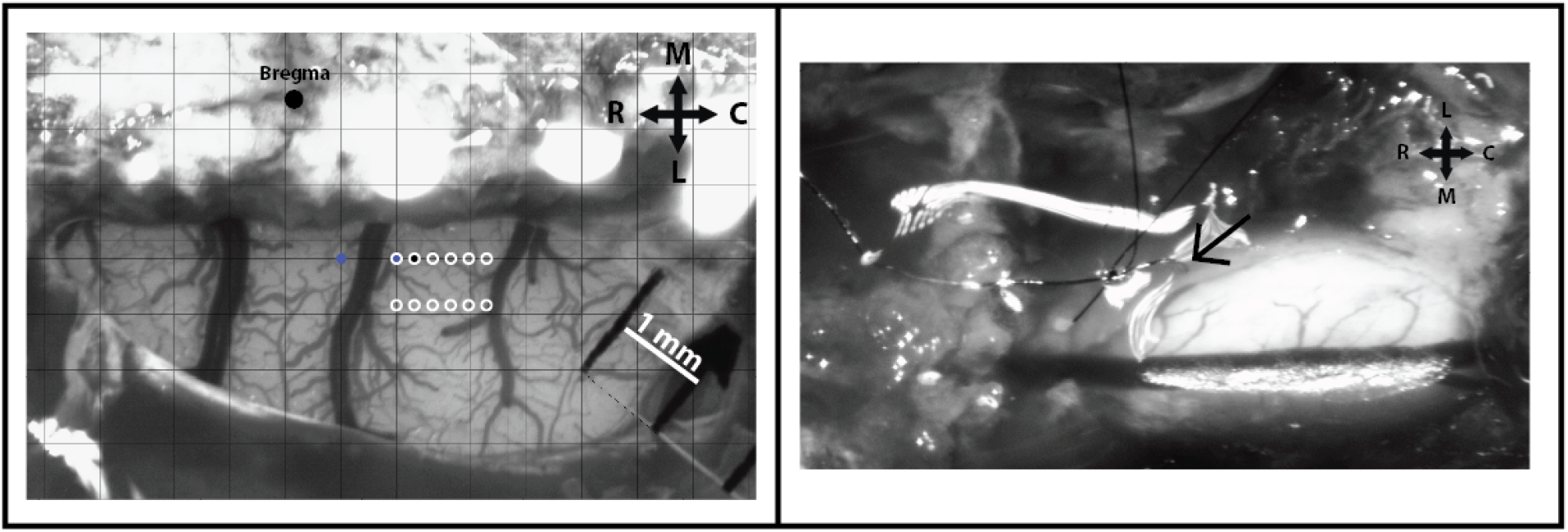
Chronic electrode implantation. **Left:** Cranial opening over hindlimb motor cortex with superimposed grid for ICMS mapping. The large black dot represents Bregma; the small blue dot represents ICMS-evoked forelimb movement; smaller black dot represents a no-response movement during ICMS; white outlined circles represent placement of chronic microarray. For this exemplar ICMS map, only two ICMS mapped sites were necessary to determine the location of the hindlimb motor cortex due to the thick vasculature of this exemplar motor map. Due to the size of the recording microarray, it was placed between the two thicker blood vessels caudal of Bregma to reduce the number of blood vessels ruptured during implantation. **Right:** Spinal cord opening over hindlimb spinal cord. The microwire was inserted into the spinal cord (black arrow) and sutured to the dura mater. For this example, only one site (black arrow) was needed to find a hip-evoked ISMS site.

To record spikes (i.e., action potentials) in the hip representation of HLA, a chronic microwire array (Tucker-Davis Technologies, Alachua, FL) was inserted into the defined area of the cortex. The recording array was a 16-channel (2 × 6 array) probe with wire diameter of 33 μm, wire length of 2 mm, wire spacing of 175 μm, row spacing of 500 μm, and tip angle of 60 degrees. Using a Kopf hydraulic microdrive (Kopf Instruments, Tujunga, CA), the array was inserted into the cortex so that electrodes were positioned at a depth of ~1700 μm, approximating Layer V of the cerebral cortex. Kwik-Cast (World Precision Instruments Inc, Sarasota, FL), a silicone elastomer, was applied directly to the cortex around the inserted wires of the recording electrode to act as a sealant. The ground wire of the recording electrode was attached to the rostral-most screw, and the impedance of each channel was recorded at the time of implantation. The average impedance values (mean ± SE) at the time of implantation were calculated by averaging all recording channels of each recording array (ADS group = 41.3 ± 3.4 kΩ; Control group = 31.7 ± 3.6 kΩ; *p* = 0.1088).

For placement of the stimulating electrode in the spinal cord, a laminectomy was performed on the T13 vertebrae similar to previous studies (Borrell et al., 2020a; Borrell et al., 2020b). The dura mater was left intact to minimize cerebrospinal fluid leakage. A custom-made fine-wire stimulating electrode was fabricated from 30-μm-diameter platinum/iridium (80%/20%) wire insulated with a 4 μm layer of polyimide (California Fine Wire, Grover Beach, CA) (Bamford et al., 2017). At least four microwires were soldered onto a custom-made electrode interface board that was soldered to an 18-pin male nano dual row Omnetics connector with guidepost holes (Omnetics Connector Corporation, Minneapolis, MN). The interface board was encapsulated in UV-cured glue to ensure stability and electrical isolation of each wire. The ground wire was wrapped around the same screw as the ground wire from the recording electrode. The microwires, not including the ground wire, were routed into a 1-mm diameter polyethylene tube (A-M Systems, Inc., Sequim, WA) to protect the wires from damage during implantation. Microwire tips were de-insulated for a length of 1 mm by mechanically removing the polyimide. Then, the tips were cutting at a 45° angle leaving 30—60 μm of exposed metal from the tip. The tip ends were then gently bent to a 90° angle with a length of ~2.27 mm from tip to bend.

The microwire implantation procedure in the hindlimb spinal cord followed the procedure developed by Bamford, *et. al.* (Bamford et al., 2017). The microwire implant was sterilized using ethylene oxide (EtO) gas. The Omnetics connector (i.e., headstage) of the microwire implant was positioned on the right hemisphere of the skull next to the recording electrode (Figure 2). The polyethylene tube and microwires were tunneled subcutaneously to the hindlimb spinal cord. Dental acrylic was used to anchor the recording and stimulating electrodes to the skull and screws. Once the acrylic was dry, the polyethylene tube of microwires was sutured to the T12 vertebral process with 6-0 polypropylene suture (PROLENE^®^ blue monofilament, taper point, C-1; 13 mm 3/8c; Ethicon) and fixed further with a small amount of surgical glue (3M Vetbond Tissue Adhesive, St. Paul, MN) being careful to glue the polyethylene tube directly to the bone and not the surrounding tissue. Before microwire insertion, a small hole was made in the dura mater using the tip of a 30-gauge hypodermic needle. Microwire insertion began by gently holding the wire with fine forceps near the tip. While avoiding major blood vessels, the tip then penetrated the superficial dorsal layers of the cord, and the remaining portion of the wire was inserted vertically until reaching the 90° bend (~2.27 mm of wire which reaches the ventral horn of the rat hindlimb spinal cord). The inserted microwire can be seen in Figure 3 Right. Only one microwire was used for the study, and when the implanted wire was deemed functional the remaining three microwires were removed.

The target for microwire placement in the spinal cord was a hip site, allowing functional pairing between the recording electrode in cortex and the stimulating electrode in spinal cord. The rostro-caudal placement of the microwire in the hindlimb spinal cord was guided by previously derived ISMS-evoked movement maps in rat (Borrell et al., 2017). The medio-lateral placement of the microwire was on the right side of the spinal cord, approximately 0.8—1.0 mm from the central blood vessel. A low-amplitude current (one biphasic pulse with 0.2 ms square-wave duration delivered at 300 Hz) was used to test the evoked movement. The current was increased until a visible movement was produced, and the threshold current intensity was recorded. If the ISMS-evoked movement was not a hip movement, the wire was removed and reinserted in another rostro-caudal location depending on the evoked movement. Every additional insertion of the microwire increases tissue damage; therefore, the number of reinsertions was limited to a maximum of three for any given location. If the maximum number reinsertions was reached and no hip movements were produced, the microwire was left in place, and the rat was deemed a control rat (i.e., same injury, surgical procedures, chronic implants, and behavioral recordings with no stimulation therapy). The microwire was laid flat on the epidural surface and were sutured with an 8-0 nylon suture (black monofilament, taper point, BV130-5, 6.5 mm 3/8c, Ethicon) directly to the dura mater. Typically, the 8-0 suture was driven through the dura mater before microwire implantation to avoid pulling the microwire out of the spinal cord. A drop of surgical glue was applied to the point of insertion and Kwik-Sil (World Precision Instruments Inc, Sarasota, FL) was used to adhere the microwire to the epidural surface and to act as a sealant. A layer of thin plastic film made of polyethylene, cold sterilized in 100% Ethyl Alcohol and allowed to dry, was measured to the size of the exposed laminectomy and used to cover the implanted region of the spinal cord. The film was affixed to the vertebral edges of the opening with small drops of surgical glue to prevent the invasion of connective tissue into the laminectomy site during recovery. Muscle layers were then sutured over the opening before suturing the skin incision closed. The skin around the headstages on the skull was pulled tight around the acrylic cap and sutured to create a snug fit. The impedance of the microwire electrode was recorded at the time of implantation. The average impedance values (mean ± SE) at the time of implantation were calculated by averaging all microwire electrodes (ADS group = 50.4 ± 8.3 kΩ; Control group = 39.0 ± 18.5 kΩ; *p* = 0.6125). The rats were allowed to recover for a few days before the beginning of ADS therapy to avoid dislodging of the microwire implants.

### ADS Paradigm

During ADS therapy, electrode leads/connectors of the headstages were tethered to the TDT neurophysiological recording and stimulation system (Tucker-Davis Technologies, Alachua, FL). Rats were placed in a 30-cm × 30-cm × 52-cm Plexiglas chamber during each ADS session. Before each ADS session, a test stimulus was applied to measure movement threshold for ISMS. A custom-made code was created using the TDT system to administer ADS. Individual spikes were detected, discriminated, and sorted in real time using principal component analysis. A consistent spiking profile was chosen from the site in the hindlimb motor cortex (i.e., the channel of the recording array that is closest to the hip representation site determined during ICMS mapping) and used to trigger stimulation in the paired site in the ventral horn of the hindlimb spinal cord. The ISMS stimulus consisted of a single, biphasic pulse (200 μs per phase; cathodal leading) from an electrically isolated stimulation circuit. Stimulation amplitude was set at 50% movement threshold determined during the implantation surgery to mitigate interference with the rat’s motor performance or muscle fatigue. A time delay of 10 ms between recorded cortical spike and triggered spinal cord stimulation pulse was used because this delay resulted in enhanced ICMS-evoked spinal cord responses in ADS sessions conducted in anesthetized SCI rats (Borrell et al., 2020b). A blanking interval of 28 ms following each spike discrimination was used to reduce the possibility of producing a positive-feedback loop, in which spinal stimulation might drive action potentials in HLA, retriggering stimulation of spinal cord (Guggenmos et al., 2013). Each ADS session occurred for 4 hours/day, 4 days/week, for 4 weeks (Figure 2).

### BBB Behavioral Assessment

The Basso, Beattie, and Bresnahan (BBB) locomotor rating scale was used as the primary behavioral outcome, as it is a sensitive measure of locomotor ability after SCI (Basso et al., 1995). Briefly, rats ran along a straight alley with the home cage and bedding at the end to encourage the rats to transverse the open field. Two examiners observed each rat to minimize any bias and to reduce the risk of missing behaviors. Scores were recorded for the left and right sides of each animal. If SCI rats did not exhibit a deficit post-injury or did not have a BBB score between 13—15 at 4-weeks post-SCI, they were removed from the study.

### Horizontal Ladder Rung Walking Test

To quantify skilled locomotor movements, rats were trained on the horizontal ladder rung walking test apparatus (Otto Environmental, Greenfield WI). This is considered to be a sensitive test to show deficits in corticospinal tract connectivity (Metz and Whishaw, 2002). The horizontal ladder consisted of side walls made of clear Plexiglas and a metal rung (3 mm diameter) walkway (Figure 4). The metal rungs could be inserted to create a floor with a minimum distance of 1 cm between the rungs. The side walls were 1 m long and 19 cm high measured from the height of the rungs, while the width of the alley was adjusted to 1 cm wider than the rat to prevent the animal from turning around. The ladder was elevated well above the ground with two refuges at each end. The starting refuge was an open container, and the ending refuge was a darkened safe box (goal box). Two patterns were used during this experiment to prevent the rats from learning the pattern and to modify the difficulty of the task. Pattern A consisted of a regular pattern with the rungs spaced at 2 cm intervals, while Pattern B consisted of an irregular pattern with the rungs spaced between 1 and 3 cm. All rats were recorded crossing the horizontal ladder five times per session for each pattern.

**Figure 4.**
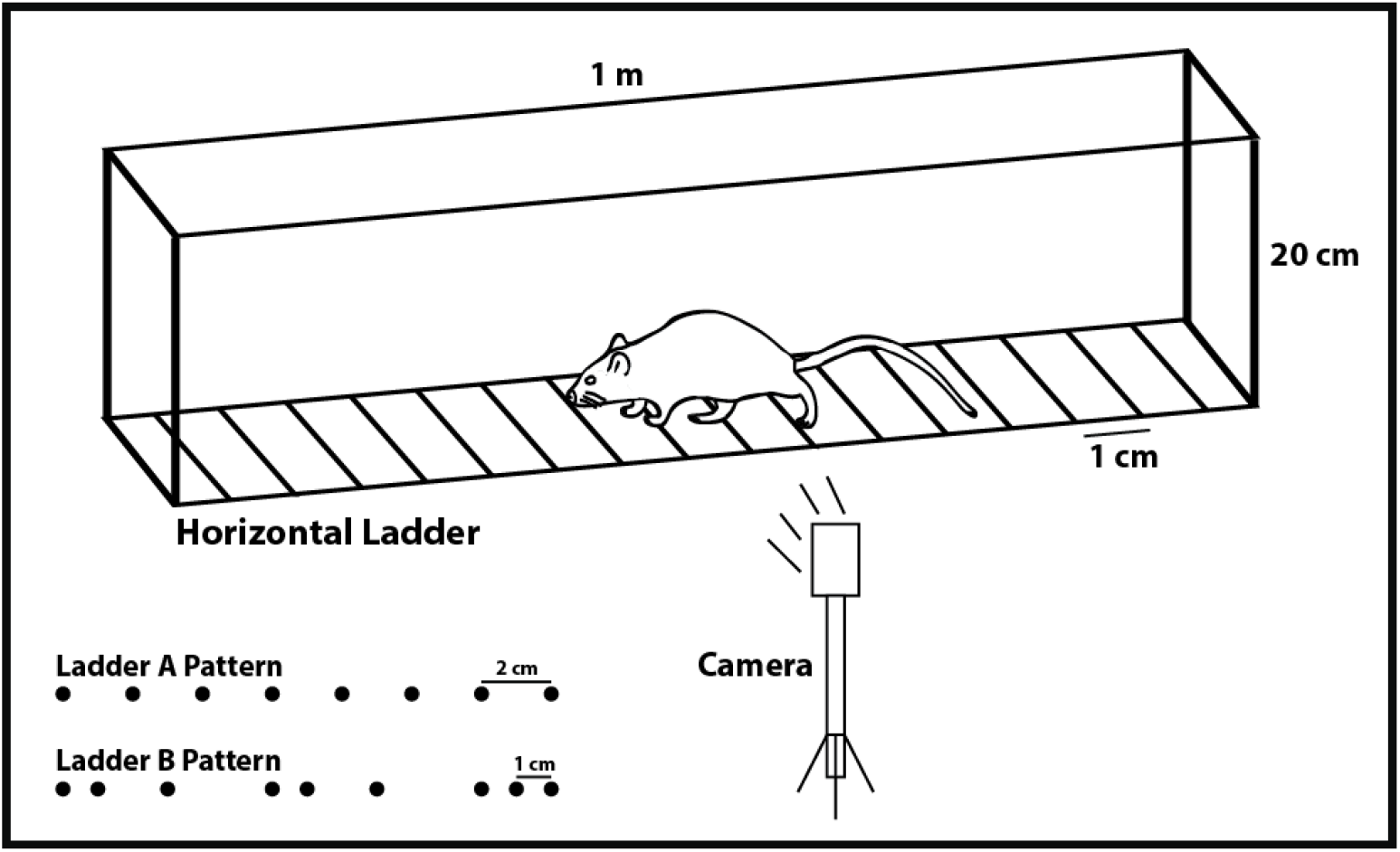
Horizontal Ladder Set-up. The camera captures the foot-faults of the paws as the rat transverses the horizontal ladder.

Rats were recorded traversing the horizontal ladder using a Sony digital camera. The camera was positioned at a slight angle so that both right and left paw positions could be recorded simultaneously. The video recordings were scored using frame-by-frame analysis at 30 frames/sec and scored and analyzed using procedures previously described (Metz and Whishaw, 2002). Only consecutive steps of each hindlimb were analyzed, so if the rat stopped, the last step before the stop and the first step after the stop were not scored. The last stepping cycle performed at the end of the ladder was not included in the scoring. The types of foot and paw placement on the rungs were rated using a 7-category scale. (0) Total miss: 0 points were given when the limb completely missed the rung and a fall occurred. (1) Deep slip: 1 point was given when the limb was placed on the rung and then slipped off when weight-bearing which caused a fall. (2) Slight slip: 2 points were given when the limb was placed on a rung, slipped off when weight bearing, but did not result in a fall and interrupt the walking. (3) Replacement: 3 points were given when the limb was placed on a rung, but before it was weight bearing, it was quickly lifted and placed on another rung. (4) Correction: 4 points were given when the limb aimed for one rung but was then placed on another rung without touching the first one. (5) Partial placement: 5 points were given when the limb was placed on the rung with either heel or digits of a hindlimb. (6) Correct placement: 6 points were given when the midportion of the palm of the limb was placed on a rung with full weight support. If different errors occurred at the same time, the lowest of the scores was recorded. Error rate in stepping was recorded as a percentage of the number of misplaced steps divided by the total number of steps on the ladder [100 × (steps with error/total steps taken)]. Steps with error corresponded to values of 0 – 3 in the foot fault scoring system.

### Tapered/Ledged Beam Test

To assess deficits in balance and fine motor control, rats were trained on the tapered/ledged beam test. The ledged beam is a highly sensitive test that allows chronic deficits to be easily scored. More specifically, the ledge of the beam provides a place to step with the impaired limbs so that the rats are not induced to compensate using alternative motor strategies (Shallert et al., 2002). The beam-walking apparatus consisted of a tapered beam with underhanging ledges (2 cm wide) on each side to permit foot faults without falling. The end of the beam was connected to darkened safe (goal) box (20.5 cm × 25 cm × 25 cm). Rats had to cross the elevated tapered beam over 60 cm in length. The width of the beam began at 6.0 cm and narrowed to a width of 1.5 cm at the end. The rats were videotaped using a Sony digital camera, and steps for each hindlimb were scored as a full-slip or a half-slip if the limb touched the side of the beam. Steps onto the ledge were scored as a full-slip and a half-slip was given if the limb touched the side of the beam. The slip ratio of each hindlimb (number of slips/number of total steps) was later calculated and averaged over three trials. The mean of three trials was used for statistical analysis.

### OptiTrack Recording

Hindlimb kinematics were recorded using a high-speed OptiTrack motion capture system (NaturalPoint, Inc., Corvallis, OR), combining six Flex 3 cameras (100 Hz; three cameras per hindlimb). Rats traversed an open field for 6 recordings/trials. A minimum of 10 step cycles was extracted for each hindlimb (excluding the first and last two steps of each trial) during this task. A total of 12 reflective markers (6 markers per hindlimb) were attached bilaterally overlying anatomical landmarks of the hindlimbs. Markers were placed on the iliac crest, greater trochanter (hip), lateral femoral epicondyle (knee), lateral malleolus (ankle), fifth metatarsophalangeal joint (mtp), and distal end of the fourth digit (digit). The 3D position of the markers was reconstructed offline using Motive Optical motion capture software. Joint angles and kinematic analysis were calculated via custom-written code in Matlab (The Mathworks, Inc., Natick, MA). Specifically, stance time, stride time, stride frequency, stride length, step height, hip range of motion (ROM), knee ROM, ankle ROM, and metatarsal ROM were the main parameters of interest with the OptiTrack. These gait parameters quantify deficits and recovery during specific phases of the over-ground gait cycle. Similar parameters have been used by other investigators for this purpose (DiGiovanna et al., 2016).

### DigiGait Recording

Gait analysis was performed with the Mouse Specifics DigiGait System (Mouse Specifics, Inc., Quincy, MA). Computerized digital footprint images were generated by high-speed video recording from the ventral aspect of the animal. A predetermined speed of 11 cm/sec was used and remained consistent throughout the experiment. All gait dynamic indices, over 40 metrics, were calculated with version 12.1 of the Mouse Specifics analysis software. Specifically, paw angle, propel duration, swing duration, stance duration, stride length, paw area, and paw overlap were the main parameters of interest on the DigiGait. These gait parameters quantify deficits and recovery during specific phases of the gait cycle. Similar parameters have been similarly used by other investigators for this purpose in rats (Krizsan-Agbas et al., 2014).

### Perfusion & Histology

At the conclusion of the study (after Week 8), rats were euthanized with an intraperitoneal injection of sodium pentobarbital (Beuthanasia-D; 100 mg/kg), then transcardially perfused with fixative (4% paraformaldehyde in 0.1 M PBS) and the spinal cord was removed. The spinal cord was transversely sectioned at 30 μm on a cryostat and sections stained with cresyl violet for histological verification of the lesion and microwire placement.

Verification that the fine-wire spinal cord implant remained in the spinal cord throughout the study was made during the dissection and removal of the spinal cord implant. Additionally, histological post-mortem examination of the lumbar spinal cord demonstrates an observable microwire path in the cord (Figure 5).

**Figure 5.**
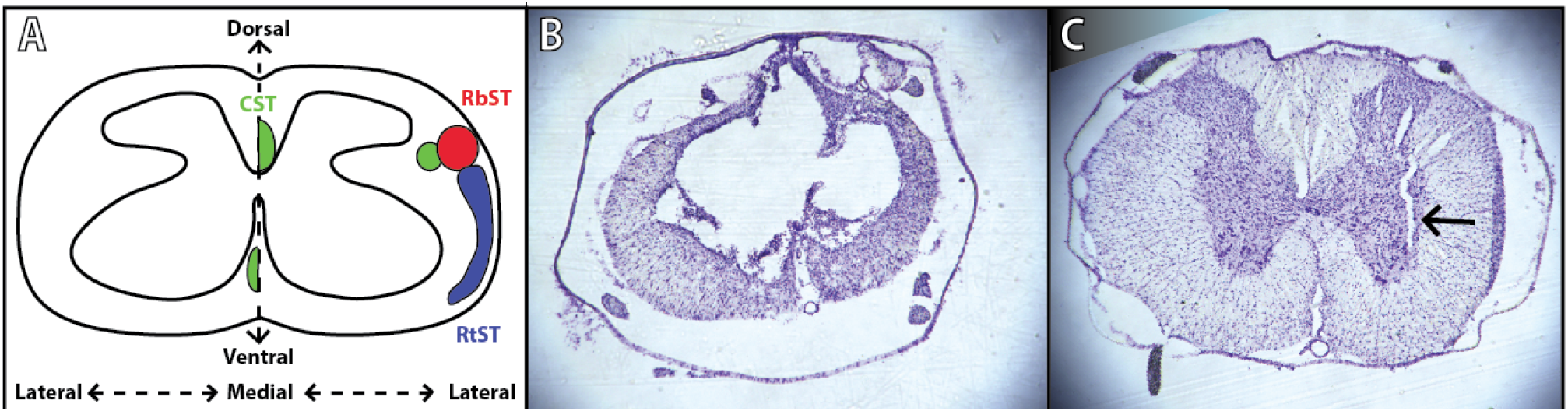
Verification of spinal cord injury and microwire implantation. **A)** Schematic diagram of a coronal thoracic section identifying the location of the descending corticospinal tract (CST; green), rubrospinal tract (RbST; red), and reticulospinal tract (RtST; blue) (Lemon, 2008; Fink and Cafferty, 2016). **B)** Representative image of spinal cord injury epicenter under T8 vertebrae stained with cresyl violet 8 weeks after injury. **C)** Representative image of microwire implant under T13 vertebrae stained with cresyl violet 4 weeks after implantation.

### Statistical Analysis

Statistical analyses of motor performance measures were conducted using JMP 11 software (SAS Institute Inc., Cary, NC). For parametric data, a two-factor repeated measures analysis of variance (ANOVA) was used to analyze effects of ADS therapy on hindlimb function over time on the skilled-movement behavioral tasks (Ledged Beam and Horizontal Ladder tasks). Hindlimb performance was obtained as a percentage of foot-faults on the behavioral tasks. An arcsin transformation was used to normalize percentage data for parametric analysis. Group comparisons were made following a significant interaction with Fisher’s Least Significance Difference (Fisher’s LSD) test. Kinematic data was also analyzed using a two-factor repeated measures ANOVA followed by group comparisons at each time point with Fisher’s LSD. The nonparametric, Mann-Whitney U test was used to compare ordinal-scaled BBB scores between the ADS and Control group. Bonferroni correction for multiple comparisons was used to control for multiple comparisons. Parametric data are presented as mean +/− standard error of the mean. Non-parametric data are presented as medians.

## Results

### SCI and Microwire Implant Verification

The average impact displacement was 1069.67 ± 31.38 μm for the ADS group and 1096.00 ± 37.92 μm for the Control group, with a total average displacement of 1082.83 ± 23.80 μm (mean ± SE) for all rats. The impact displacement value was not significantly different between groups. Exemplar transverse histological sections through the center of the injury and at the location of the microwire tract are shown in Figure 5. After SCI, the spinal grey matter at the epicenter was severely damaged. The ventral and lateral spinal cord white matter tracts (the location of the ventral CST and RtST) remained largely intact, while the dorsal column white matter tracts (location of the dorsomedial CST) appear to have been severely damaged (Figure 5). After microwire implantation, the microwire created a narrow dorsoventral tract which helps verify the location of the microwire (black arrow, Figure 5C). During post-mortem examination of the lumbar spinal cord, all implanted microwires from both rat groups were physically removed (pulled out with surgical tweezers) from the spinal cord. No microwire implant had become visibly dislodged throughout the study in either rat group.

### HLA Cortical Spiking & Stimulation Frequency During ADS

The average cortical spike frequency during therapy sessions for ADS rats was 8.30 ± 0.29 (mean ± SE) spikes/sec. Each recorded HLA spike was used to trigger stimulation in the spinal cord. HLA spikes were not recorded during the blanking period. Thus, HLA spike frequency is equal to stimulation frequency (hereby called spike frequency). There was no significant change in spike frequency over the course of a given four-hour ADS session. The recorded spike frequencies were further calculated for each day of ADS (Figure 6). The average recorded spike frequencies were not statistically different (*p* > 0.05) across days of ADS therapy.

**Figure 6.**
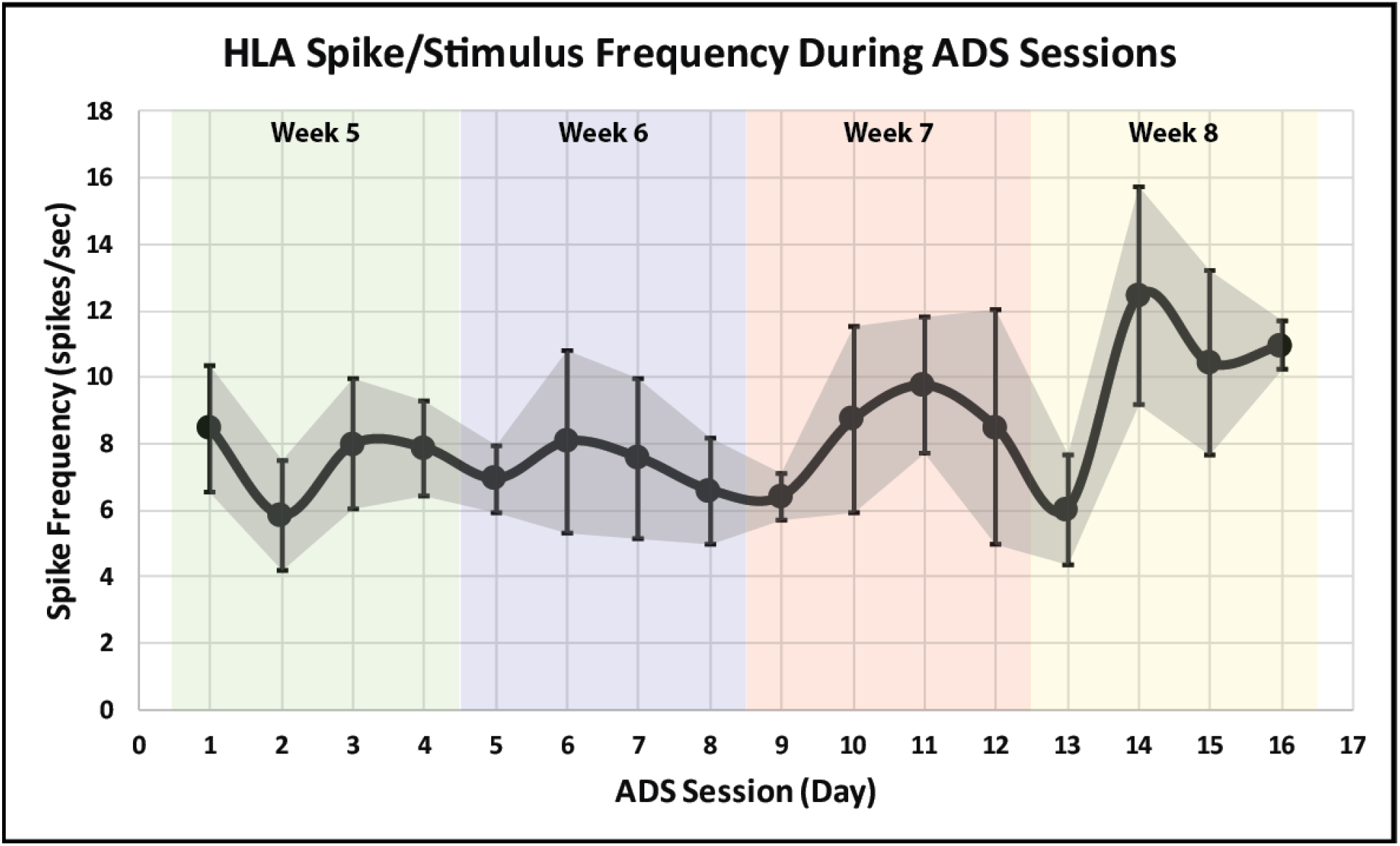
Average HLA spike/stimulus frequency (mean ± SE) during each day of ADS for the ADS rat group. The average is shown in black with error bars and grey shading representing standard error of the mean.

### ISMS Current During ADS

The minimum current needed to evoke movement (i.e., movement threshold) via the implanted microwire in the hindlimb spinal cord was recorded directly after the implantation surgery and after each week of ADS. After the implantation surgery, the average current for both groups recorded at movement threshold was 36.0 ± 7.3 μA (mean ± SE). The average current recorded at movement threshold immediately after the implantation surgery was 24.0 ± 4.3 μA for the ADS group and 55.5 ± 15.6 μA for the Control group (*p* = 0.0802). After each week of ADS, we were unable to evoke a movement via the implanted microwire in the hindlimb spinal cord. As a result, the threshold used during ADS for each rat receiving ADS was equal to the minimum current required to evoke movement as measured after implantation surgery. Although movement could not be evoked post-implant, a qualitative difference was observed between the rats in the ADS group and the Control group: Rats in the ADS group would actively stretch the right hindlimb or keep the right hindlimb completely extended. This only occurred when the stimulation current was turned on. This did not occur in the Control rats where the stimulation current remained off during therapy sessions.

### BBB Scores

#### Baseline BBB scores

Before SCI contusion, all rats were tested to verify that there were no pre-injury hindlimb deficits. All rats had a BBB score of 21 for both hindlimbs (Figures 7A and 7B) which consisted of consistent plantar stepping, consistent forelimb-hindlimb coordination during gait, consistent toe clearance, parallel paw position throughout the entire gait cycle, consistent trunk stability, and tail consistently up.

**Figure 7.**
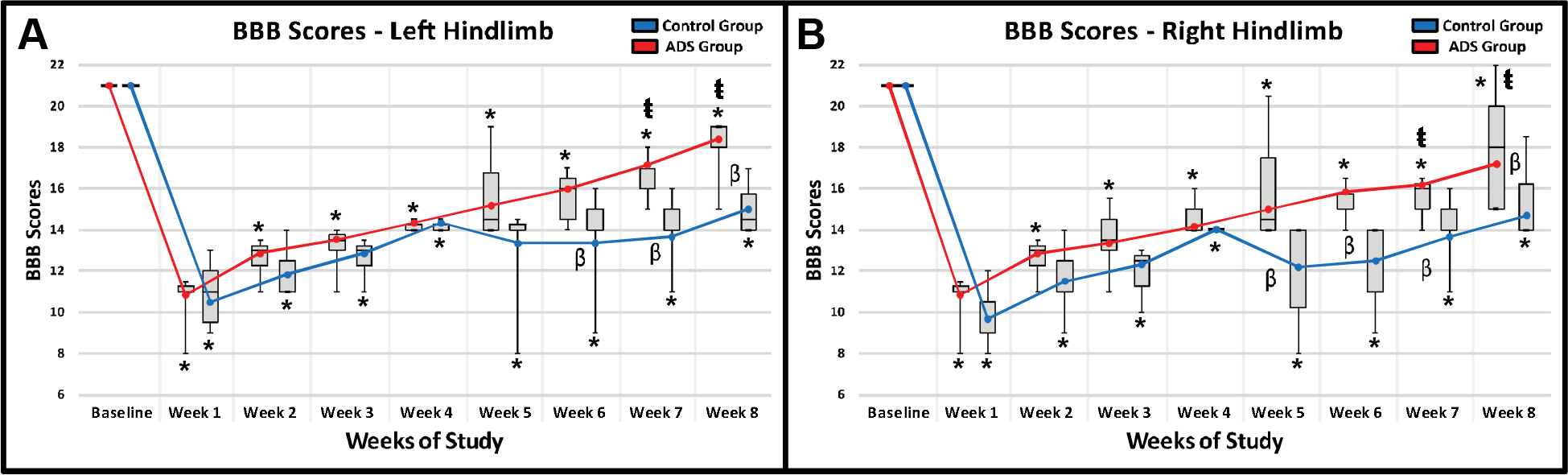
Average BBB scores (mean ± SE; line graph) and median BBB scores (median and quartiles; box and whisker plot) in ADS and Control rats. **A)** Average and median BBB scores of the left hindlimb in ADS and Control rats. **B)** Average and median BBB scores of the right hindlimb in ADS and Control rats. Error bars are ± SE. **\*** = Significant difference relative to baseline (within-group comparison; *p* < 0.05). ŧ = Significant difference relative to Week 4 (within-group comparison; p<0.05)). β = Significant difference between groups relative to each separate week (*p* < 0.05).

#### Post-SCI BBB scores

All rats exhibited clear hindlimb deficits immediately after SCI contusion. BBB scores were not significantly different between groups for both hindlimbs (Week 1). Within-group comparisons between Week 1 and Baseline displayed a significant decrease in BBB scores (Week 1 in Figures 7A and 7B; *p* < 0.0001) for both hindlimbs in both groups. In the absence of therapy, BBB scores naturally improved in both hindlimbs for 4 weeks post-SCI. The BBB scores for both hindlimbs of each rat group remained significantly lower relative to baseline BBB scores (within-group comparisons, *p* < 0.0001). After 4 weeks post-SCI (pre-implant), all rats had a BBB score within the inclusion criterion range (BBB score of 13 – 15) (Week 4, Figures 7A and 7B); which consisted of frequent/consistent plantar stepping, frequent/consistent forelimb-hindlimb coordination, no toe clearance, externally rotated paws throughout the entire gait cycle, trunk instability, and up/down tail. BBB scores were not significantly different between the ADS and Control groups at any time point post-SCI weeks 1-4.

#### Post-implant BBB scores

At one week post-implant and after one week of ADS therapy (Week 5), the ADS group displayed improved hindlimb motor function compared to the Control group in only the right hindlimb (*p* = 0.0396, Figure 7B). At two weeks post-implant and after two weeks of ADS therapy (Week 6) the BBB scores of both hindlimbs for the ADS group were significantly higher than the BBB scores of both hindlimbs for the Control group. The significant difference between groups continued at three weeks (Week 7) and four weeks (Week 8) post-implant (Figures 7A and Figure 7B). The final BBB scores in the ADS group (i.e., BBB score of ~17 – 18) represented improvement in toe clearance and in the position of the paw during gait (either parallel throughout the entire gait cycle or parallel only when the toe returns to the ground after the swing phase).

Within-group comparisons during the ADS period (weeks 5-8) compared with Week 4 (pre-ADS) show that by Weeks 7 and 8, BBB performance in the ADS group had improved significantly compared with Week 4 (Week 7: left hindlimb, *p* = 0.0062; right hindlimb, *p* = 0.0248; Week 8: left hindlimb, *p* = 0.0107; right hindlimb, *p* = 0.0088). However, despite gradual improvement over the subsequent weeks, BBB scores of the Control group were not significantly different from their Week 4 scores.

### Horizontal Ladder Rung Test

#### Baseline foot-fault scores on horizontal ladder

Before SCI contusion, all rats demonstrated minimal foot-faults, averaging zero to two foot-faults for each hindlimb while traversing the ladder (Figures 8A and 8B).

**Figure 8.**
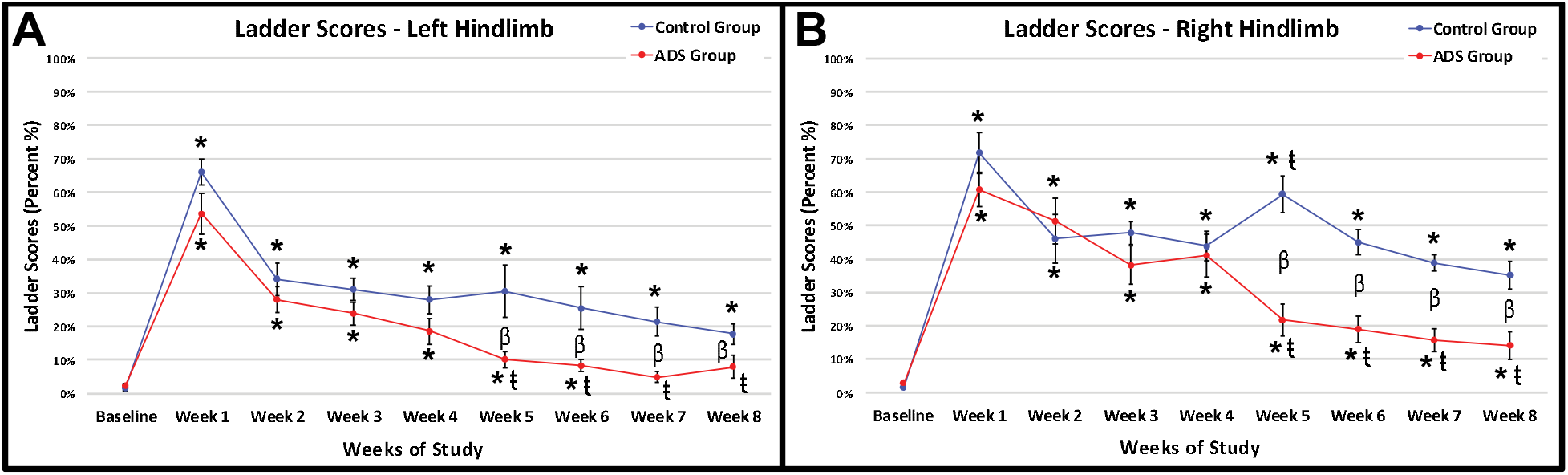
Average foot-fault scores (mean ± SE) on both horizontal ladder patterns for ADS and Control rats. **A)** Average foot-fault scores of the left hindlimb for ADS and Control rats. **B)** Average foot-fault scores of the right hindlimb for ADS and Control rats. Error bars are ± SE. **\*** = Significant difference relative to baseline (within-group comparison; *p* < 0.05). ŧ = Significant difference relative to Week 4 (within-group comparison; *p* < 0.05). β = Significant difference between groups relative to each separate week (*p* < 0.05).

#### Post-SCI foot-fault scores on horizontal ladder

In post-SCI Week 1, all rats exhibited the same hindlimb deficits on the horizontal ladder after SCI contusion (no statistical difference between groups). Within-group comparisons between Week 1 and Baseline showed a significant increase in foot-faults for both hindlimbs and in both ADS and Control groups (Figures 8A and 8B; *p* < 0.0001). Foot-faults naturally improved for both hindlimbs over 4 weeks post-SCI but remained significantly higher compared to baseline (within-group comparisons between Week 4 and Baseline; both groups, both hindlimbs, *p* < 0.0001).

#### Post-implant foot-fault scores on horizontal ladder

At one-week post-implant and after one week of ADS therapy (Week 5), the ADS group had improved scores compared to the Control group (Figures 8A and 8B). The foot-fault scores for both hindlimbs remained significantly lower in the ADS group than the Control group for the remainder of the study (Weeks 6-8).

After one-week post-implant and after one week of ADS therapy (Week 5), the foot-fault scores from the ADS group significantly improved in both hindlimbs compared to Week 4 (left hindlimb, *p* = 0.0360, Figure 8A; right hindlimb, *p* = 0.0002, Figure 8B) and continued to be significantly improved for the rest of the study (Weeks 6-8). With the exception of Week 5 in the right hindlimb (Figure 8B), significant differences in foot-fault scores were not observed within the Control group after implantation and remained unchanged by week 8.

Foot-fault scores from the left hindlimb of the ADS group returned to baseline measurements (three weeks post-implant; within-group comparison between Baseline and Week 7; left hindlimb, *p* = 0.4285, Figure 8A) and remained at a non-significant level for the remainder of the study (within-group comparison between Baseline and Week 8; left hindlimb, *p* = 0.2106, Figure 8A). This improvement was similarly seen in the right hindlimb of the ADS group; however, foot-fault scores did not return to baseline levels during any week post-implant (within-group comparison between Baseline and Week 8; left hindlimb, *p* = 0.0054, Figure 8B). This recovery in foot-fault scores was not seen from either hindlimb in the Control group.

### Tapered/Ledged Beam Test

#### Baseline foot-fault scores on the ledged beam

Before SCI contusion, all rats demonstrated minimal foot-faults on the ledged beam with each hindlimb (Figures 9A and 9B).

**Figure 9.**
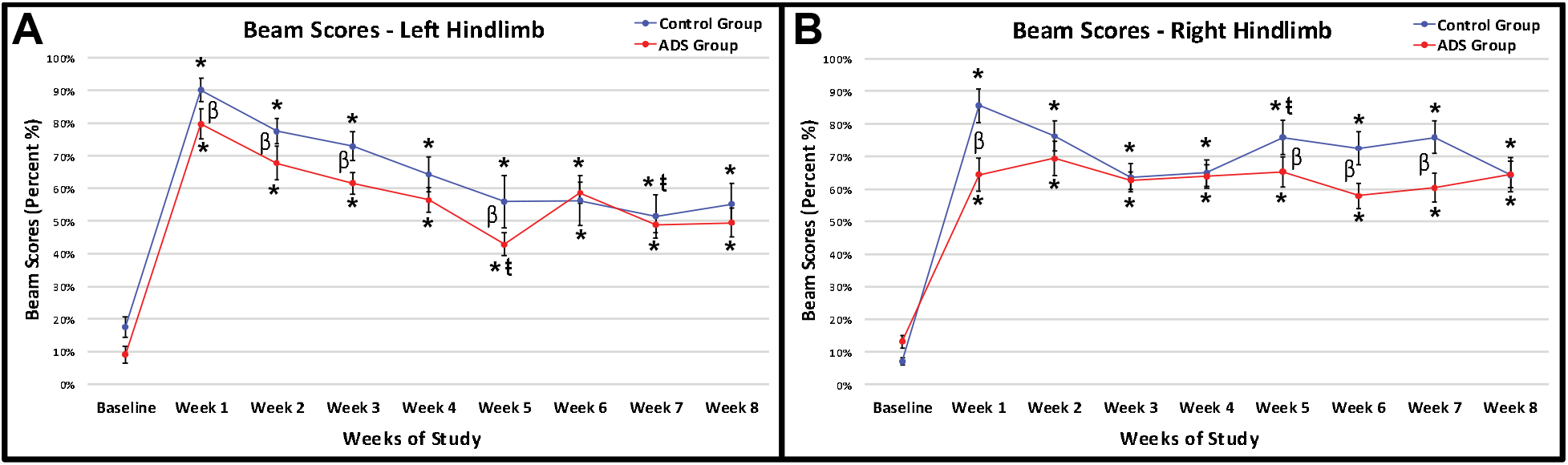
Average foot-fault scores (mean ± SE) on the tapered/ledged beam task in ADS and Control rats. **A)** Average foot-fault scores of the left hindlimb in ADS and Control rats. **B)** Average foot-fault scores of the right hindlimb in ADS and Control rats. Error bars are ± SE. **\*** = Significant difference relative to baseline (within-group comparison; *p* < 0.05). ŧ = Significant difference relative to Week 4 (within-group comparison; *p* < 0.05). β = Significant difference between groups relative to each separate week (*p* < 0.05).

#### Post-SCI foot-faults on the ledged beam

All rats exhibited hindlimb deficits on the ledged beam immediately after SCI contusion; however, there was a significant difference between groups in both hindlimbs one-week post-SCI (Week 1; left hindlimb, *p* = 0.0064, Figure 9A; right hindlimb, *p* < 0.0001, Figure 9B). Within-group comparisons between Week 1 and Baseline displayed a significant increase in foot-faults (*p* < 0.0001) for both hindlimbs in both ADS and Control groups. Foot-faults naturally improved for both hindlimbs for 4 weeks post-SCI. After 4 weeks post-SCI (i.e., pre-implant), all rats still displayed a deficit with significantly increased foot-fault scores in both hindlimbs and for both groups. Significant differences in foot-fault scores between-groups were not observed by Week 4.

#### Post-implant foot-fault scores on the ledged beam

After one-week post-implant and after one week of ADS therapy (Week 5), foot-fault scores in the ADS group began to significantly decrease for the left hindlimb on the ledged beam as compared to pre-implant foot-fault scores (within-group comparison of Weeks 4 and 5; left hindlimb, *p* = 0.0454, Figure 9A). This decrease in the left hindlimb was greater in the ADS group compared to the Control group (between-group comparison of Weeks 4 and 5; *p* = 0.0032; Figure 9A). Significant differences in foot-fault scores within-group relative to week 4 were not observed by Week 8.

### OptiTrack Results

#### Knee range of motion (ROM)

Both groups displayed clear deficits in knee ROM for both hindlimbs 1 week post-SCI. However, the knee ROM between-groups were significant weeks 2-4 before implant). After implant surgery, there was no significant changes in knee ROM of both hindlimbs for each group for the remainder of the study). Significant differences in ROM between groups were not observed by week 8.

#### Remaining OptiTrack parameters

Stance time, swing time, stride time, stride frequency, stride length, step height, hip ROM, ankle ROM, and metatarsal ROM all displayed deficits after SCI. However, most parameters showed similar between-group differences prior to ADS therapy as was seen with knee ROM. Thus, any significant differences in measurements between groups after Week 4 may not be attributable to ADS therapy.

### DigiGait Parameters

#### DigiGait parameters

Clear deficits in paw angle, propel duration, swing duration, stance duration, stride length, paw area, and paw overlap distance were measured by the DigiGait analysis after Week 1 post-SCI in both hindlimbs. These deficits remained 4 weeks post-SCI before therapy. After ADS therapy, no significant improvement was observed in any of the tested DigiGait parameters.

## Discussion

This proof-of-concept study demonstrates that activity dependent stimulation (ADS) using spike-triggered intraspinal microstimulation (ISMS) can enhance behavioral recovery of locomotor function after spinal cord injury, as measured by behavioral assessment. Previous studies have used ADS paradigms specifically focused on recovery of forelimb function (Moritz et al., 2008; Guggenmos et al., 2013; Zimmermann and Jackson, 2014; McPherson et al., 2015). Efforts have been made to recover hindlimb function using spike-triggered epidural stimulation across a unilateral spinal cord laceration (Capogrosso et al., 2016). The present study demonstrates that an ADS approach using cortically-evoked intraspinal stimulation may enhance recovery of function specific to hindlimb control. Importantly, behavioral recovery was seen after ADS (i.e., spike-triggered ISMS) was applied across a bilateral spinal cord contusion injury that more accurately mimics injuries typically seen in the clinic.

### Hindlimb Motor Deficits After Spinal Cord Injury

Hindlimb deficits after a bilateral, moderate thoracic spinal cord contusion, like the injury model used in this study, have been well reported for the BBB open field assessment. From the onset of injury, hindlimb function naturally improves until 4 weeks post-injury, when recovery plateaus (Basso et al., 1995; Scheff et al., 2003; Krizsan-Agbas et al., 2014). Differences regarding the natural recovery of hindlimb motor function and the BBB score recorded at 4 weeks post-injury have been related to the location of and severity of injury on the thoracic spinal cord. However, these reports agree that natural improvement in post-injury hindlimb function plateaus around 4 weeks post-injury.

Hindlimb deficits recorded on the DigiGait four weeks after contusion SCI in this study were similar to those reported in previous studies (Ek et al., 2010; Krizsan-Agbas et al., 2014). Deficits in kinematics seen here were also similar to that of other studies using a similar contusion model (Hamers et al., 2001; Koopmans et al., 2009). Natural recovery of hindlimb motor function recorded with DigiGait and with the OptiTrack was found to plateau around 4 weeks post-SCI, similar to BBB scores reported here and by others.

Similarly, both the ladder and beam scores generally plateaued at 4 weeks post-SCI in the control group without ADS intervention. Previous studies utilizing the horizontal ladder and narrow beam tasks have examined behavioral recovery in rats after unilateral spinal cord hemisection (Ballermann and Fouad, 2006; Arvanian et al., 2009) rather than the bilateral spinal cord contusion model used here. This limits direct comparisons to published literature.

### Hindlimb Motor Function Recovered After ADS

After 4 weeks of ADS therapy, hindlimb motor function improved in the ADS group as measured by improved BBB scores and foot-fault scores on the horizontal ladder, as compared to the Control group. Improvement in motor function did not occur in the Control group and remained plateaued for the remainder of the study. Even though we were unable to evoke movement post-implant in the spinal cord, motor function still improved in the ADS group, which indicates that stimulation therapy had a positive effect on these measured outcomes.

### Potential Motor Pathways Conditioned by ADS

Three descending motor pathways provide substantial influence on hindlimb motor function in the rat: corticospinal tract (CST), rubrospinal tract (RbST), and reticulospinal tract (RtST). Crossed corticospinal axons, traveling in the dorsal columns in rats, are severely damaged regardless of spinal cord contusion severity (Basso et al., 2002). Based on the qualitative level of white matter remaining after injury, the RbST and dorsolateral CST (dCST) in the dorsolateral funiculus and the RtST and ventromedial CST (vCST) in the ventral funiculus may be relatively intact after the moderate bilateral contusion injury used in this study (Figure 5). These intact pathways may be viable targets for strengthening of connectivity from cortex to the lumbar cord to enhance the recovery of voluntary movement.

It has been shown that hindlimb spinal cord activity can still be evoked via cortico-reticulo-spinal pathways that remain intact after CST injury (Schucht et al., 2002). Sparing RtST fibers after SCI has been shown to result in the recovery of 7 to 8 points on the BBB locomotor score, as compared to a recovery of 1 to 2 points due to spared CST fibers (Schucht et al., 2002). Significant increases in BBB scores (roughly 3—4 points) were observed in the ADS group in this study for both hindlimbs. This recovery in BBB scores suggests that ADS may have strengthened the cortico-reticulo-spinal pathways and resulted in improved control of open-field locomotion.

Control of forelimb-hindlimb (FL-HL) and left-right coordination (i.e., interlimb coordination) is known to be influenced by long descending propriospinal pathways (Frigon, 2017). The amount of propriospinal sparing is uncertain in the present study, but it has been shown that long descending propriospinal axons are almost completely lost in the cervical cord with this type of injury (Basso et al., 2002). Nonetheless, FL-HL coordination was recovered before electrode implantation (BBB score of ~14 after Week 4) and remained present after ADS therapy, suggesting that some propriospinal axons remained intact. Since propriospinal pathways are known to receive a large number of terminals from reticulospinal neurons (Mitchell et al., 2016), this suggests that coordination is strongly controlled via the RtST and further supports the notion that ADS strengthened cortico-reticulo-spinal fibers. However, if propriospinal pathways are disrupted after injury, recovery of FL-HL coordination may be associated with the number of rubrospinal axons spared after injury, especially if long descending propriospinal axon sparing is insufficient (Basso et al., 2002). In addition to a strengthened cortico-reticulo-spinal pathway due to ADS, plasticity most likely occurs with the RbST depending on the remaining number of long descending propriospinal neurons after injury.

The ladder rung walking task was used to assess skilled walking (i.e., aiming, fine-adjustment, limb coordination, and balanced weight supported stepping), which is assumed to rely more strongly on CST function (Metz and Whishaw, 2002). The narrowed beam task was used to test weight bearing and fine motor adjustments during gait as well as to remove compensatory strategies that provide balance to non-impaired limbs (Schallert et al., 2002). Surprisingly, foot fault scores on the horizontal ladder walking task returned to pre-injury measures in both hindlimbs after ADS therapy; however, no improvement was observed in either hindlimb on the narrowed beam task after ADS therapy. Improvement on the narrowed beam task was not expected, as trunk stability never returned after ADS therapy, as measured in the final BBB scores. This lack of trunk stability results in an inability to balance appropriately on the beam. However, aiming, fine-adjustment, and limb coordination did improve, as reflected on the ladder rung walking task. This suggests that there may be partial recovery of the CST, especially since the CST has been shown to play a major role in the precise control over paw placement and limb trajectory (Drew et al., 2002). Due to the extent of the injury on the descending corticospinal fibers, this seems highly unlikely and suggests that the RbST may have played a role in the recovery seen on the ladder rung walking task (Basso et al., 2002). However, a strengthened cortico-reticulospinal tract cannot be ruled out as it has the capacity to deliver information from the cortex to the spinal cord in the absence of direct CST input (Mitchell et al., 2016).

Finally, the ventral CST has been shown to sprout and parallel functional recovery after a dorsal transection in the cervical enlargement of the rat (Weidner et al., 2001). It is unknown if this sprouting occurs after a thoracic contusion. Nonetheless, it is assumed that the ventral CST remains intact after injury in this study, in addition to some sparing of the dorsolateral CST. The recovery seen after ADS therapy could involve plasticity of the ventral and dorsolateral CSTs in chorus with rubrospinal and reticulospinal fibers.

### Limitations of Study

The main limitation to this study was the inability to evoke motor movement 1—3 days post-implant. This could be due to one or a combination of the following factors: 1) Extensive scar tissue formed around the microwire thus driving the maximum current needed to evoke movement (i.e., movement threshold) well above the maximum current that was used in this study (i.e., 100 μA with 1 biphasic pulse); 2) The microwire shifted in the spinal cord but was not removed from the spinal cord. It is uncertain from the histological cross-section of the microwire tract if the microwire shifted in the spinal cord; however, the narrow medio-lateral width of the tract would suggest that it remained relatively stationary. Additionally, the recorded impendences of the microwire throughout the study indicate that we were able to drive the correct current during ADS and that scarring around the implanted microwire was minimal. Nonetheless, there may have been enough scarring around the microwire to inhibit the ability to evoke movement. Finally, the behavioral recovery seen after each week of ADS suggests that the microwire was still able to produce a neuromodulatory effect that aided in the observed improved measurements of behavioral recovery.

Additionally, the fine-wire electrode that was implanted in the right side of the spinal cord may have itself caused deficits, as noted by instances of decreased performance measures in the Control group in the first tests post-implant. These deficits were noted in some rats one week after implantation (Week 5) as a decrease in BBB scores and an increase in foot-faults on the Horizontal Ladder and the Ledged Beam tasks. However, this initial deficit was not observed in any of the rats in the ADS group. This could be an indication that stimulation after implantation provides some benefit that offsets a deficit caused by the implant. However, since behavioral tasks were not conducted immediately after implantation surgery and only after ADS therapy began, it is uncertain if the rats in the ADS group obtained a similar deficit due to the implant that was observed in the Control group. In order to address this issue, future studies will need to conduct behavioral tasks immediately after electrode implantation and before any stimulation therapy begins.

## Conclusion

Spike-triggered intraspinal microstimulation was shown to significantly improve hindlimb motor performance after a moderate spinal cord contusion as measured by improved BBB scores and horizontal ladder performance. The behavioral recovery shown here indicates that cortically-driven activity dependent stimulation may have strengthened cortico-reticulospinal fibers in parallel with plasticity of rubrospinal, dorsolateral corticospinal, and ventral corticospinal fibers. These data will inform the future development of activity-dependent based therapies which have been shown to be translatable across mono- and multi-synaptic pathways as well as various injuries related to the central nervous system.

## Acknowledgments

This work was supported by the Paralyzed Veterans of America Research Foundation #3068, The Ronald D. Deffenbaugh Family Foundation, NIH/NINDS R01 NS030853, T32 Neurological Rehabilitation Sciences Training Program, and NIH/NINDS F31 NS105442. The authors thank Erica Hoover, Brad Lamb, and Matthew Jaeschke for exceptional technical support.

